# The superficial white matter in language processing: Broca’s area connections are bilaterally associated with individual performance in children and adults

**DOI:** 10.1101/2025.07.31.666959

**Authors:** Shiva Hassanzadeh-Behbahani, Zhou Lan, Leo Zekelman, R. Jarrett Rushmore, Yuqian Chen, Tengfei Xue, Suheyla Cetin-Karayumak, Steve Pieper, Yanmei Tie, Edward Yeterian, Alexandra J. Golby, Nikos Makris, Fan Zhang, Yogesh Rathi, Lauren J. O’Donnell

**Affiliations:** Department of Radiology, Brigham and Women’s Hospital, Harvard Medical School, Boston, Massachusetts, USA; Center for Clinical Investigation, Channing Division of Network Medicine, Brigham and Women’s Hospital, Boston, Massachusetts, USA; Department of Neurosurgery, Brigham and Women’s Hospital, Harvard Medical School, Boston, Massachusetts, USA; Speech and Hearing Bioscience and Technology, Harvard University, Cambridge, Massachusetts, USA; Department of Psychiatry, Brigham and Women’s Hospital, Harvard Medical School, Boston, Massachusetts, USA; Department of Anatomy and Neurobiology, Chobanian and Avedisian School of Medicine, Boston University, Boston, Massachusetts, USA; Center for Morphometric Analysis, Departments of Psychiatry and Neurology, A. Martinos Center for Biomedical Imaging, Massachusetts General Hospital, Charlestown, Massachusetts, USA; Isomics, Inc., Cambridge, Massachusetts, USA; Department of Psychology, Colby College, Waterville, Maine, USA; Harvard-MIT Health Sciences and Technology, Massachusetts Institute of Technology, Cambridge, Massachusetts, USA

**Author notes:** **Corresponding author information:** Lauren J. O’Donnell and Shiva Hassanzadeh-Behbahani.

**Keywords:** Adolescent Brain Cognitive Development Study, Human Connectome Project Young Adult, superficial white matter, tractography, language

## Abstract

While deeper white matter connections, such as the arcuate fasciculus and frontal aslant tract, are well known for their role in language and show leftward asymmetries in adults, the contribution of the short-range cortico-cortical superficial white matter (SWM) connections remains less understood. In this preregistered study, we examined white matter connections of Broca’s area and its right hemisphere homolog in early adolescents and young adults using two large, publicly available datasets: the Adolescent Brain Cognitive Development Study and the Human Connectome Project Young Adult Study, totaling over 10,000 participants. We anatomically curated the O’Donnell Research Group (ORG) tractography atlas to identify SWM fiber clusters intersecting Broca’s area (pars opercularis and pars triangularis), confirmed through expert visual inspection. We investigated the microstructure, structural connectivity, and lateralization of Broca’s area SWM and its relationship with language performance (Picture Vocabulary and Oral Reading Recognition assessments), in comparison to the deeper white matter connections of the frontal aslant tract and arcuate fasciculus. The arcuate fasciculus demonstrated the strongest and most consistent leftward lateralization of both microstructure (fractional anisotropy, FA) and structural connectivity (number of streamlines, NoS), with structure-function associations that were bilateral in adolescents and left-dominant in adults. Interestingly, despite weaker lateralization than the arcuate fasciculus, both the SWM and the frontal aslant tract demonstrated comparable associations with language performance. The frontal aslant tract showed greater leftward lateralization (FA and NoS) with age and was bilaterally associated with language performance, particularly in adolescents. Compared to these deeper tracts, Broca’s area SWM demonstrated left-lateralized FA and right-lateralized NoS in both adolescents and adults, with stronger FA lateralization in adults. Bilateral SWM FA and NoS were significantly associated with language performance in both hemispheres and age groups. Overall, these results suggest that Broca’s area SWM may support language in a more bilaterally distributed manner and highlight the importance of considering SWM connections in studies of language development and neurosurgical planning.

## 1. Introduction

The white matter neuroanatomy of the language system has been focused principally on the long association fiber tracts connecting the language-related areas of the frontal, temporal, and parietal areas. More specifically, Broca’s area in the frontal lobe, Wernicke’s area in the temporal lobe, and the inferior parietal lobule were considered to be interconnected mainly by the arcuate fascicle (e.g., (Catani & Mesulam, 2008; Dejerine & Dejerine-Klumpke, 1895; Geschwind, 1970). Although this connectional network was originally thought to involve only the dominant hemisphere for language, more recently this view has been revised and broadened to include cortical regions in the non-dominant hemisphere as well (e.g., (Hickok & Poeppel, 2004, 2007)). Furthermore and more recently, a “superior to inferior frontal gyrus pathway connection” (Lawes et al., 2008) or frontal aslant tract (FAT) (Catani et al., 2012, 2013) has been added to the list of fiber connections related to language. The FAT has been considered as a medium-range or type 2 fiber tract connection by our group (Makris et al., 2023) and as a medium association fiber bundle (Van Dyken et al., 2024). Despite these advances in understanding language-related white matter tracts, there remains a significant paucity of reports related to the specific involvement of the superficial white matter (SWM) in language processing (e.g., (Duffau et al., 2005; Ford et al., 2010; Oishi et al., 2008)).

The SWM, composed of short-range cortico-cortical association fibers situated between the deep white matter and the cortex (Oishi et al., 2011), plays an important role in cognition, neurodevelopment, aging, and disease (Van Dyken et al., 2024). These local connections, which facilitate communication between nearby gyri, account for up to 60% of total white matter volume, roughly twice that of the long-range fiber bundles traditionally emphasized in diffusion MRI research (Schüz & Braitenberg, 2002). Recent studies increasingly demonstrate the feasibility of mapping the SWM with diffusion MRI tractography (Guevara et al., 2020; K. Schilling et al., 2025; K. G. Schilling et al., 2023; Van Dyken et al., 2024; Xue et al., 2023; F. Zhang et al., 2024). While recent tractography studies support the role of the SWM in human cognitive function (Lo et al., 2025; Wang et al., 2025), its specific contribution to language remains largely unexplored.

Broca’s area, classically defined as the pars opercularis and pars triangularis of the left inferior frontal gyrus (Brodmann areas 44 and 45), is crucial for speech production and language processing (Flinker et al., 2015; Papoutsi et al., 2009; Schnur et al., 2009). Diffusion MRI studies consistently show leftward asymmetries in major deep white matter tracts connected to Broca’s area, including the arcuate fasciculus and FAT (Lebel & Beaulieu, 2009; Linn et al., 2024; Sreedharan et al., 2015). A few small studies have also reported asymmetries in SWM microstructure, particularly in frontal regions (Phillips et al., 2013; Román et al., 2022).

The present study characterizes the anatomy and hemispheric lateralization of Broca’s area SWM, compares its features with those of deeper language-related pathways of the arcuate fasciculus and frontal aslant tract, and examines how SWM microstructure and connectivity relate to language performance in early adolescents and young adults. We draw on two large, publicly available diffusion MRI datasets: the Adolescent Brain Cognitive Development (ABCD) Study (Cetin-Karayumak et al., 2024; Jernigan et al., 2018) and the Human Connectome Project Young Adult (HCP-YA) study (Elam et al., 2021; Van Essen et al., 2013), which together include over 10,000 participants spanning early adolescence and young adulthood. By extending our anatomically curated tractography atlas (F. Zhang et al., 2018), we extract and analyze both deep and superficial white matter connections of Broca’s area (Figure 1). We preregistered this study by documenting our research plan prior to data analysis on the Open Science Framework (https://osf.io/j45k7).

**Figure 1.**
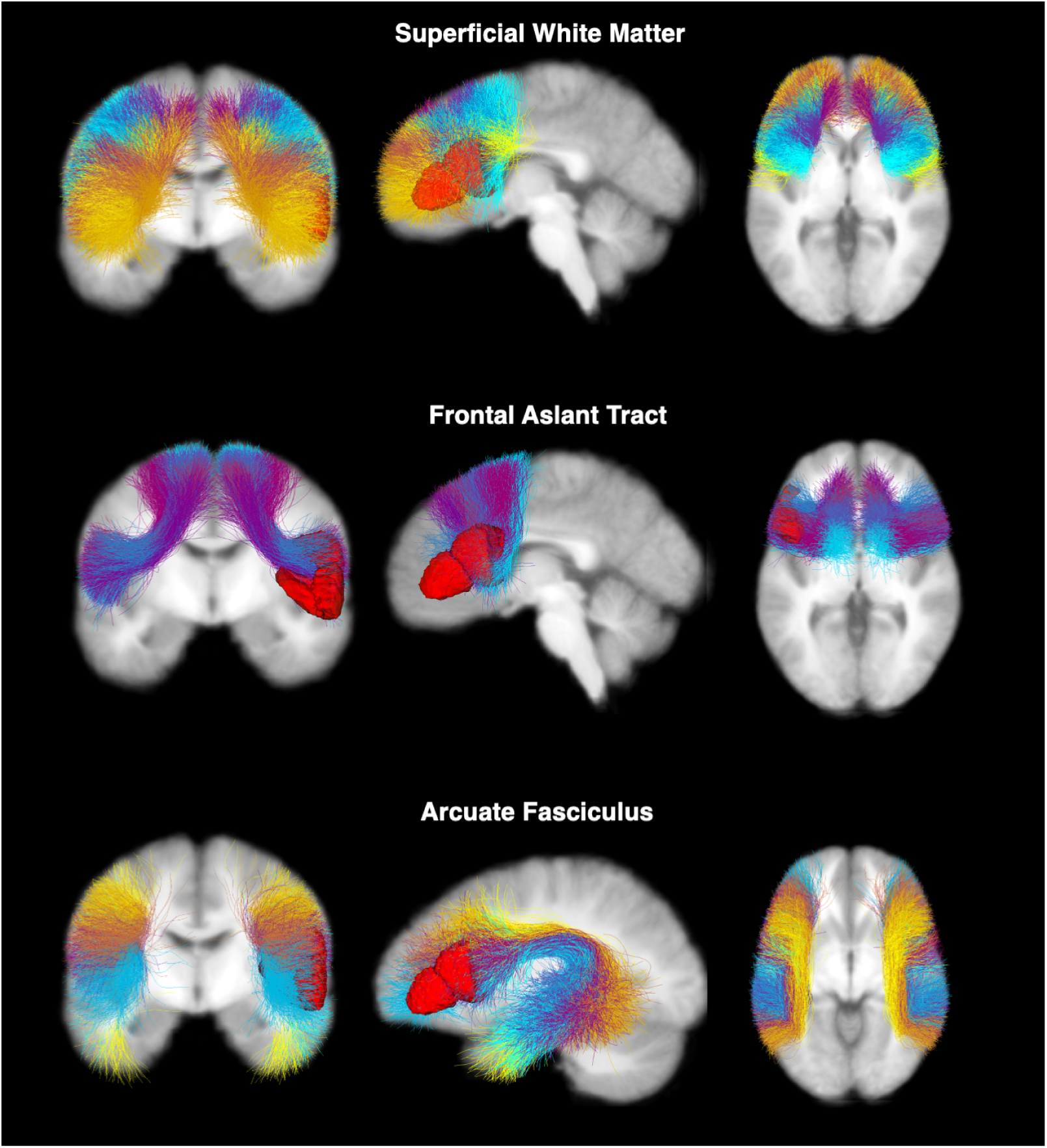
White matter connections of Broca’s area from the ORG atlas, shown as anatomically curated fiber clusters. Anterior, left, and superior views show Broca’s area superficial white matter, frontal aslant tract, and arcuate fasciculus. 2D images (located behind the 3D connections) are provided for visual reference. Broca’s area is displayed in the left hemisphere as a red 3D surface model, and it is defined as pars opercularis and pars triangularis of the left inferior frontal gyrus using the FreeSurfer parcellation. Fourteen SWM fiber clusters (shown in different colors) connecting Broca’s area and different regions of the frontal cortex are identified and combined for statistical analyses. For comparison, we also study the FAT and arcuate fasciculus.

## 2. Methods

We investigate the microstructure, connectivity, and lateralization of Broca’s area connections, leveraging diffusion MRI tractography and an atlas-based machine learning method for the identification of deep and superficial white matter connections. In this section, we describe the datasets and processing (Section 2.1), the anatomical curation of the atlas (Section 2.2), and the statistical analysis (Section 2.3).

### 2.1 Datasets

We utilized previously computed diffusion MRI tractography data and language performance scores from two large datasets, as follows.

#### 2.1.1 Participants

We employed two large, publicly available datasets: the ABCD Study of early adolescents (n=9,345; 4,464 females; 8,282 right-handed; ages 8.9-11, mean age 9.9 ± 0.6) (Cetin-Karayumak et al., 2024) and the HCP-YA study of young adults (n=1,065; 575 females; 966 right-handed; ages 22-37, mean age 28.75 ± 3.67), totaling over 10,000 participants. All participants provided informed consent prior to inclusion in the ABCD or HCP-YA studies (Clark et al., 2018). The ABCD Study received centralized IRB approval from the University of California, San Diego, with local IRB approvals obtained at each participating site (Auchter et al., 2018). The HCP-YA study was approved by the Washington University IRB.

#### 2.1.2 Language performance data

The ABCD and HCP-YA datasets provided two language performance assessments from the NIH Toolbox Cognition Battery: the Picture Vocabulary Test (PicVocab) and the Oral Reading Recognition Test (ReadEng) (Gershon et al., 2014). These assessments measured individual performance on two different aspects of language processing. PicVocab is an assessment of vocabulary knowledge, and ReadEng is an assessment of reading decoding skills (Gershon et al., 2014). Higher scores on these assessments reflect better vocabulary abilities or reading skills.

#### 2.1.3 Diffusion MRI acquisition and processing

ABCD dMRI scans were acquired with multiband EPI (slice acceleration factor=3, 1.7 mm isotropic resolution) with 96 directions, seven b=0 volumes, four b-values (b=500, 1000, 2000, 3000 s/mm²) with 6, 15, 15, and 60 directions respectively, and TR/TE 4100/88 ms (Siemens) and 4100/81.9 ms (GE) (Hagler et al., 2019). HCP-YA dMRI scans were acquired with multiband EPI (slice acceleration factor=3, 1.25 mm isotropic resolution) with 270 directions, eighteen b=0 volumes, and three b-values (b= 1000, 2000, 3000 s/mm²) with 90 directions each, collected with TR/TE=5520/89.5 ms. Both datasets underwent minimal pre-processing (eddy current, motion, and distortion correction, and isotropic resampling) (Glasser et al., 2013; Hagler et al., 2019).

Because ABCD scans were acquired across 21 sites, we used the harmonized version of the dMRI data, which corrects for scanner-, site-, and protocol-related biases using a rotation-invariant spherical harmonics approach that preserves inter-subject variability (Cetin Karayumak et al., 2019; Cetin-Karayumak et al., 2024). The HCP-YA dataset required no harmonization, as all scans were acquired on a single scanner (Van Essen et al., 2012).

Details of the dMRI tractography processing and parcellation procedures have been described previously (Cetin-Karayumak et al., 2024; Zekelman et al., 2022). From the minimally pre-processed HCP and harmonized ABCD dMRI data, the b=3000 shell and all b=0 volumes were selected, as the b=3000 shell is optimal for resolving crossing fibers and represents the most comparable acquisition between ABCD and HCP-YA (Cetin-Karayumak et al., 2024; Descoteaux et al., 2007; Ning et al., 2015; Zekelman et al., 2021). Tractography was performed using the Unscented Kalman Filter (UKF) tractography method (Malcolm et al., 2010; Reddy & Rathi, 2016), a multi-tensor approach implemented in ukftractography^1^ that provides fiber-specific measures of tissue microstructure such as fractional anisotropy (FA). Whole-brain tractography in the ABCD and HCP-YA datasets was then parcellated (Cetin-Karayumak et al., 2024; Zekelman et al., 2022) using the whitematteranalysis^2^ (WMA) machine learning approach. WMA consistently parcellates tractography across the lifespan, health conditions, and acquisitions with high test-retest reliability (F. Zhang et al., 2018, 2019). WMA employs a neuroanatomically curated pathway atlas, the O’Donnell Research Group (ORG) atlas (F. Zhang et al., 2018), to identify large white matter connections and finely parcellated fiber clusters. Multiple groups have employed the ORG atlas to study SWM fiber clusters in health and disease (Guevara et al., 2020; Wang et al., 2025; Xue et al., 2023; D. Zhang et al., 2024; F. Zhang et al., 2018).

### 2.2 Anatomical curation of Broca’s area connections

The fiber clusters in the ORG atlas were previously curated by a neuroanatomist into recognized pathways, including major deep white matter tracts such as the arcuate fasciculus, as well as a category of SWM fiber clusters (F. Zhang et al., 2018).

We performed further neuroanatomical curation of the ORG atlas to identify fiber clusters intersecting Broca’s area, including the SWM, FAT, and arcuate fasciculus. Broca’s area was defined as the pars opercularis and pars triangularis based on the Desikan-Killiany Atlas cerebral cortical parcellation implemented in FreeSurfer (Desikan et al., 2006). We used the groupwise anatomical parcellation provided with the ORG atlas to obtain the anatomical labels of the FreeSurfer regions intersected by each fiber cluster. Each fiber cluster under study was visually inspected by the neuroanatomist team (RJR, EY, NM).

First, we identified 14 SWM fiber clusters connecting to Broca’s area. Next, we delineated the FAT to include 7 fiber clusters connecting Broca’s area and the superior frontal gyrus, in line with established neuroanatomical definitions of the FAT. Clusters were visually evaluated by neuroanatomists (NM, RJR, EY). Finally, we identified 11 arcuate fasciculus fiber clusters from the ORG atlas; of these, 8 were found to connect to Broca’s area and were the focus of our analysis. One male ABCD participant was excluded from analyses as the right hemisphere arcuate fasciculus was not detected.

### 2.3 Quantification, statistical analysis, and hypotheses tested

For each individual’s SWM, FAT, and arcuate fasciculus connections, we studied dMRI measures of fractional anisotropy (FA) and number of streamlines (NoS), which are widely used to study brain microstructure, cognition, and development (F. Zhang et al., 2022). These quantitative measures were previously estimated for all subjects in the ABCD and HCP-YA datasets (Cetin-Karayumak et al., 2024; Zekelman et al., 2022). The fiber clusters belonging to each pathway of interest were aggregated into a single structure for statistical analysis. FA was averaged across all streamline points in a pathway, while NoS was the total number of streamlines in a pathway (F. Zhang et al., 2022). We computed the hemisphere laterality index (LI) of FA and NoS using the following formula (Banfi et al., 2019; Thiebaut de Schotten et al., 2011): (*right value − left value*)/(*right value + left value*). Accordingly, LI values range from −1 to 1, with negative values indicating left-hemisphere lateralization and positive values indicating right-hemisphere lateralization. The absolute value of LI reflects the strength of lateralization, with values closer to 1 indicating stronger lateralization.

To evaluate whether white matter connections were significantly lateralized, we tested whether LI values differed significantly from zero, following established best practices (Banfi et al., 2019; Broce et al., 2015; Thiebaut de Schotten et al., 2011). We first adjusted LI values for age and sex using multiple linear regression models, with LI as the dependent variable and age and sex as independent variables. We then conducted one-sample t-tests on the adjusted LI values to assess whether their mean significantly differed from zero, indicating population-level lateralization (Banfi et al., 2019). These analyses were performed separately in the ABCD and HCP-YA datasets, using LI values derived from both FA and NoS values. To compare lateralization between early adolescents and young adults, we conducted two-sample t-tests.

To evaluate associations between white matter microstructure (i.e., FA) and language performance, we conducted multiple linear regression analyses with the left or right white matter FA as the dependent variable and PicVocab or ReadEng assessment scores, age, and sex as independent variables. White matter structural connectivity (NoS) can be considered as count data in regression analysis (Peterson, 1999). Following the state-of-the-art statistical methodology for count data analysis, we applied negative binomial regression (Hilbe, 2011), a popular model that models count variables and handles over-dispersion, to evaluate associations between NoS and language performance. Specifically, we conducted negative binomial regression analyses with the left or right white matter NoS as the dependent variable and PicVocab or ReadEng assessment scores, age, and sex as independent variables.

In addition, to evaluate associations between white matter lateralization and language performance, we used multiple linear regression analyses with the LI of a white matter connection as the dependent variable and PicVocab or ReadEng assessment scores, age, and sex as independent variables.

In all analyses, we used the standard *p* < .05 criteria to test for statistical significance. We corrected for multiple comparisons using the false discovery rate (FDR) (Benjamini & Hochberg, 1995), applied separately within each dataset. For regression models evaluating the relationship between white matter properties and language performance, FDR correction was applied separately for FA and NoS, with 12 comparisons each. For regressions assessing LI and language performance, FDR correction was applied separately for LI of FA and LI of NoS (6 comparisons each). For t-tests assessing LI, FDR correction was applied across six tests. All analyses were performed in R statistical software (R Core Team, 2021). Broca’s area white matter visualizations were created in 3D Slicer with SlicerDMRI (Norton et al., 2017).

## 3. Results

### 3.1 Curation and identification of Broca’s area white matter connections

Figure 2 provides a visualization of the SWM, FAT, and arcuate fasciculus connections in the atlas and in one example participant from each dataset. We identified 7 fiber clusters belonging to the FAT and 14 Broca’s area SWM fiber clusters connecting to Brodmann areas 44 and 45. Visualizations of individual SWM fiber clusters in the atlas and in one example participant from the ABCD and HCP-YA datasets are provided in Figures 3-5.

**Figure 2.**
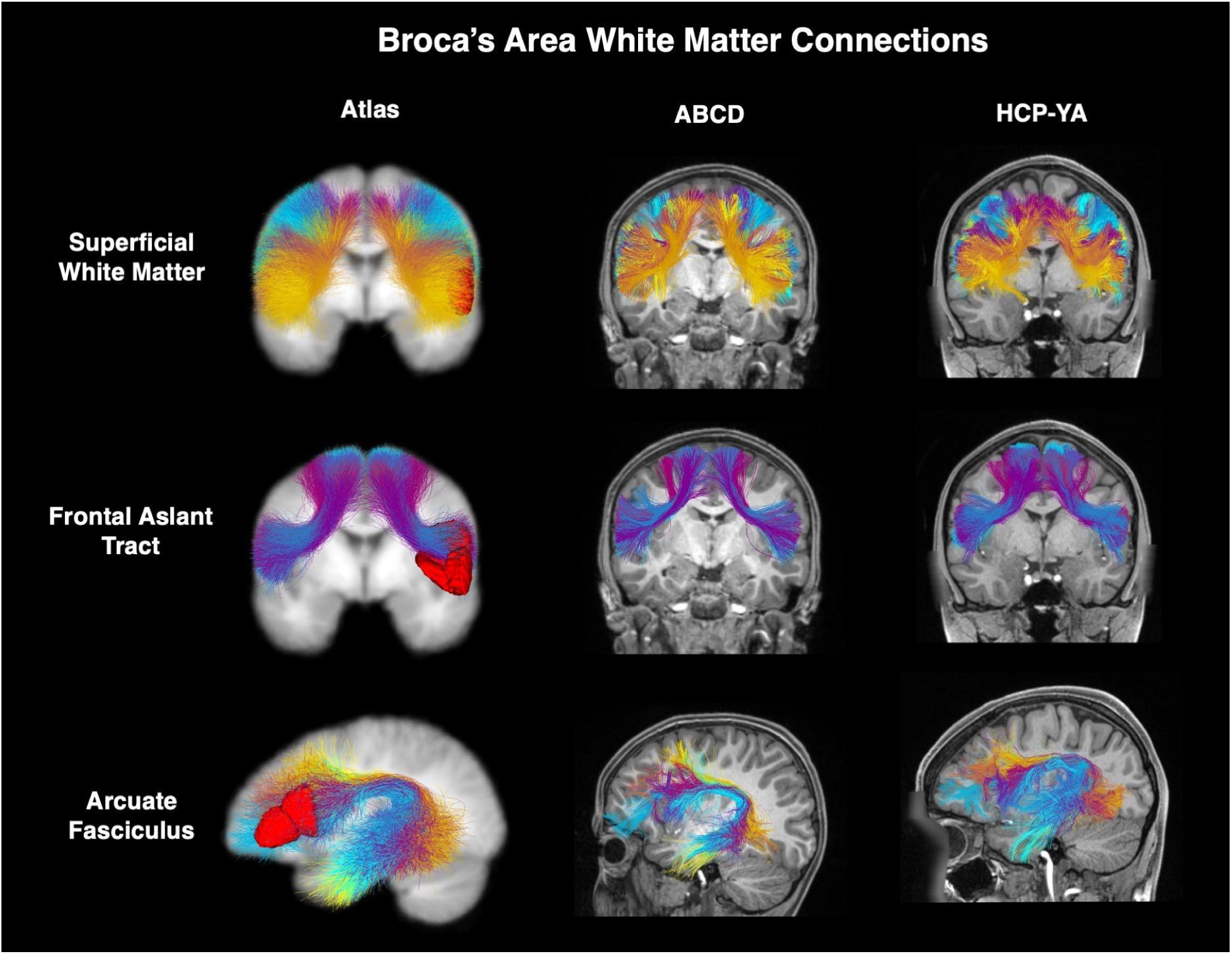
Broca’s area white matter connections. SWM, FAT, and arcuate fasciculus connections are visualized in the ORG atlas and in an individual ABCD and HCP-YA participant. 2D coronal/sagittal image planes (located behind the 3D connections) are provided for visual reference. Broca’s area is displayed in the left hemisphere as a red 3D surface model.

**Figure 3.**
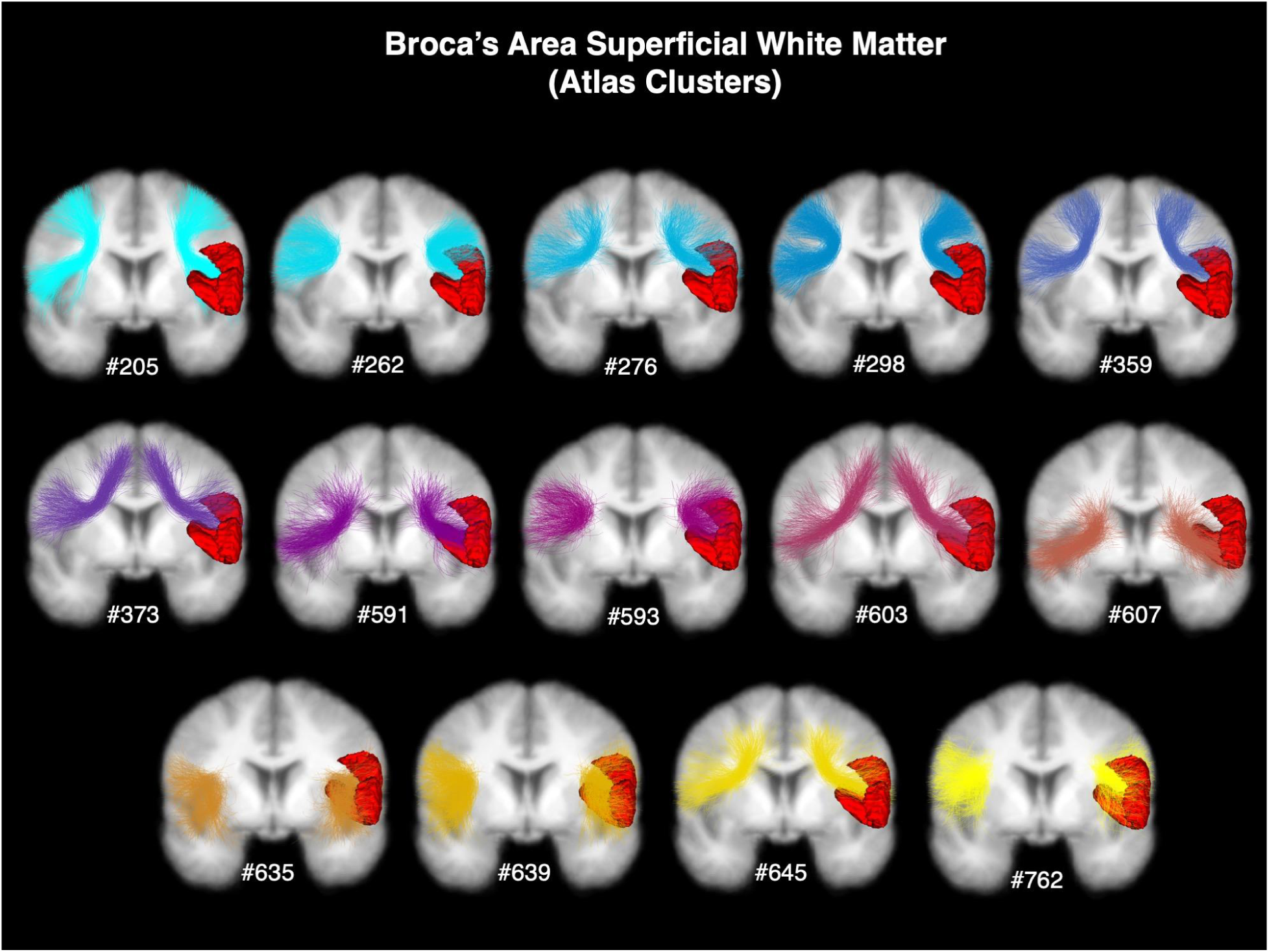
Visualizations of individual Broca’s area SWM fiber clusters in the ORG atlas. Numbers correspond to the ORG atlas fiber cluster number. Broca’s area is displayed in the left hemisphere as a red 3D surface model. Also see Figure 1 for additional views. 2D images (located behind the 3D connections) are provided for visual reference.

**Figure 4.**
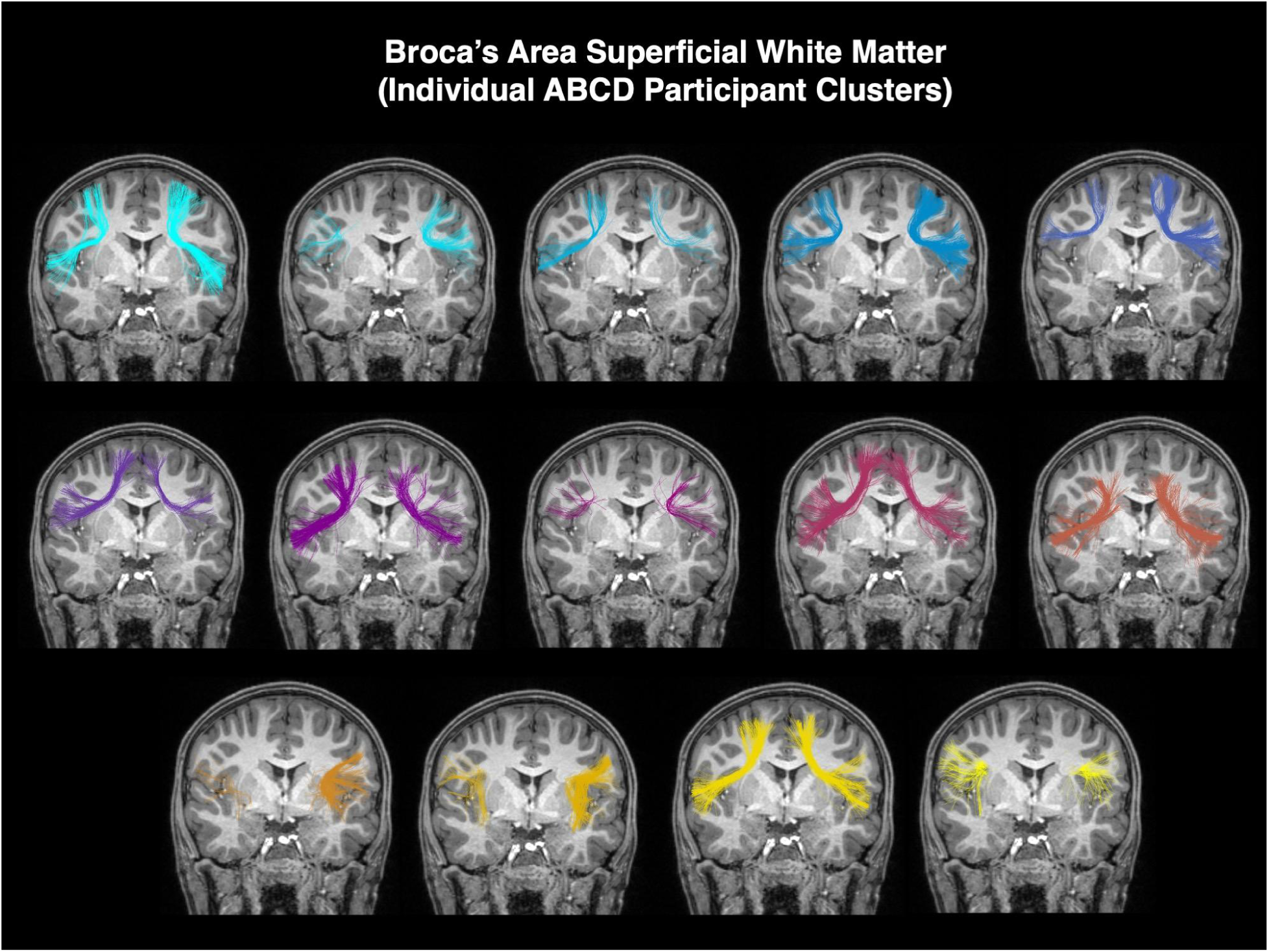
Visualizations of individual Broca’s area SWM fiber clusters in one example participant from the ABCD dataset. 2D images (located behind the 3D connections) are provided for visual reference.

**Figure 5.**
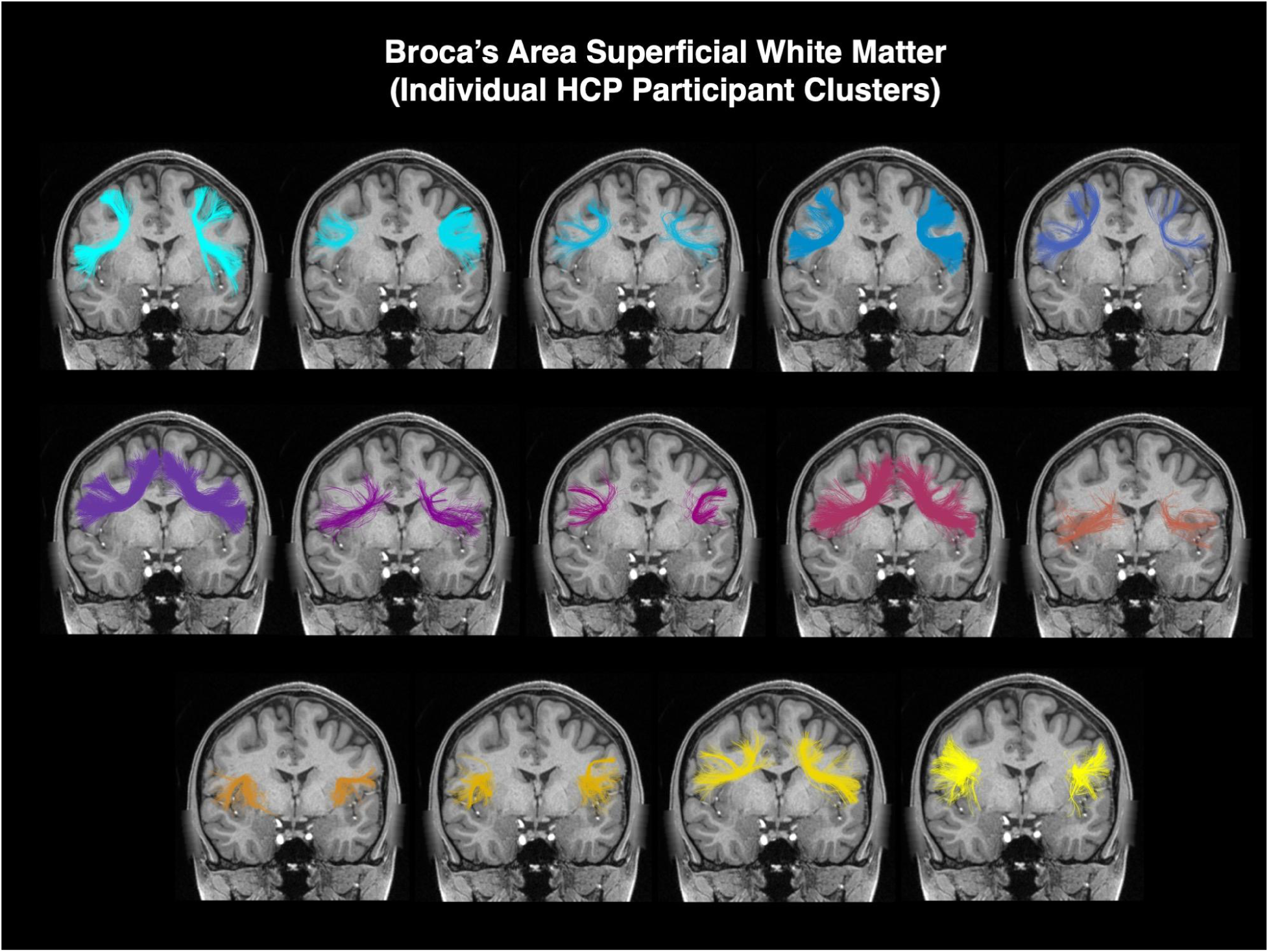
Visualizations of individual Broca’s area SWM fiber clusters in one example participant from the HCP-YA dataset. 2D images (located behind the 3D connections) are provided for visual reference.

### 3.2 Structural lateralization of Broca’s area superficial white matter

Figure 6 provides the LI values (Figure 6A) and distribution of FA and NoS (Figure 6B) in the early adolescent (ABCD) and young adult (HCP-YA) datasets.

**Figure 6.**
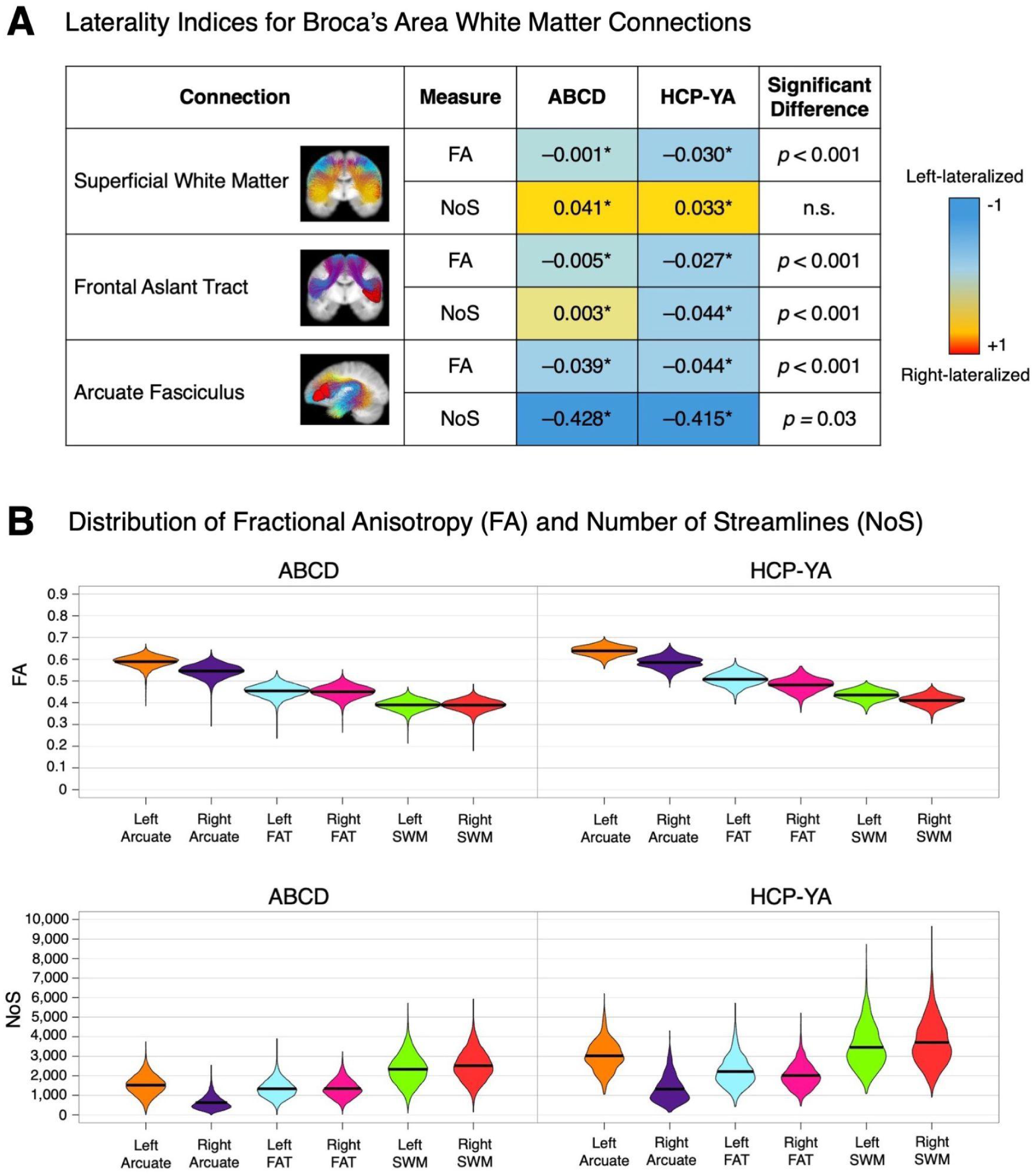
Quantitative measures of Broca’s area connections in early adolescents (ABCD) and young adults (HCP-YA). (A) Laterality indices calculated from FA and NoS values across datasets. Asterisks (*) indicate LI values significantly different from zero (*p*<0.001, FDR-corrected). The “Significant Difference” column displays the FDR-corrected p-values assessing whether LI values differ significantly between the ABCD and HCP-YA datasets. (B) Violin plots depict the distributions of FA and NoS for each dataset; black bars represent the mean values.

In both datasets, Broca’s area SWM exhibits significant left lateralization of FA and right lateralization of NoS (all *p*<0.001). The left lateralization of SWM FA is significantly greater in young adults than adolescents (*t*(1400.3) = 37.52, *p*<0.001), while the rightward lateralization of NoS of Broca’s area SWM is consistent between the datasets.

In adolescents (ABCD), the FAT is generally symmetric, exhibiting very small magnitude but significant left lateralization of FA and right lateralization of NoS (all *p*<0.001). In young adults (HCP-YA), the FAT exhibits a larger magnitude significant left lateralization of FA and NoS (all *p*<0.001). The leftward lateralization of both FAT NoS and FA is significantly greater in young adults than adolescents (NoS: *t*(1490.6) = 8.08, *p*<0.001; FA: *t*(1396.6) = 28.08, *p*<0.001).

Consistent with prior findings (Lebel & Beaulieu, 2009), the arcuate fasciculus is left-lateralized in both FA and NoS in both early adolescents and young adults (all *p*<0.001). Leftward lateralization of the arcuate fasciculus FA is significantly greater in young adults compared to adolescents (*t*(1355.6) = 7.54, *p*<0.001). However, leftward lateralization of the arcuate fasciculus NoS is significantly greater in adolescents than in young adults (*t*(1385) = −2.26, *p*=0.03).

Of note, in both adolescents and young adults, the arcuate fasciculus lateralization is more pronounced than that of the FAT or SWM. The NoS of the arcuate fasciculus exhibits the strongest leftward lateralization (ABCD LI: –0.428; HCP-YA LI: –0.415), whereas all other laterality indices—including those of the SWM—are of much smaller magnitude.

### 3.3 Superficial white matter microstructure and connectivity relate to language performance bilaterally in early adolescents and young adults

Next, we assess relationships between Broca’s area white matter FA and NoS and language performance in early adolescents and young adults.

Multiple regression analyses of tissue microstructure (FA) reveal that FA of Broca’s area SWM in both hemispheres is positively associated with performance on language assessments (PicVocab and ReadEng) for early adolescents and young adults (all *p*<0.005; Figure 7A). Similarly, negative binomial regression analyses reveal that greater structural connectivity (NoS) in Broca’s area SWM in both hemispheres is significantly associated with better language performance for early adolescents and young adults (all *p*<0.001; Figure 7B).

**Figure 7.**
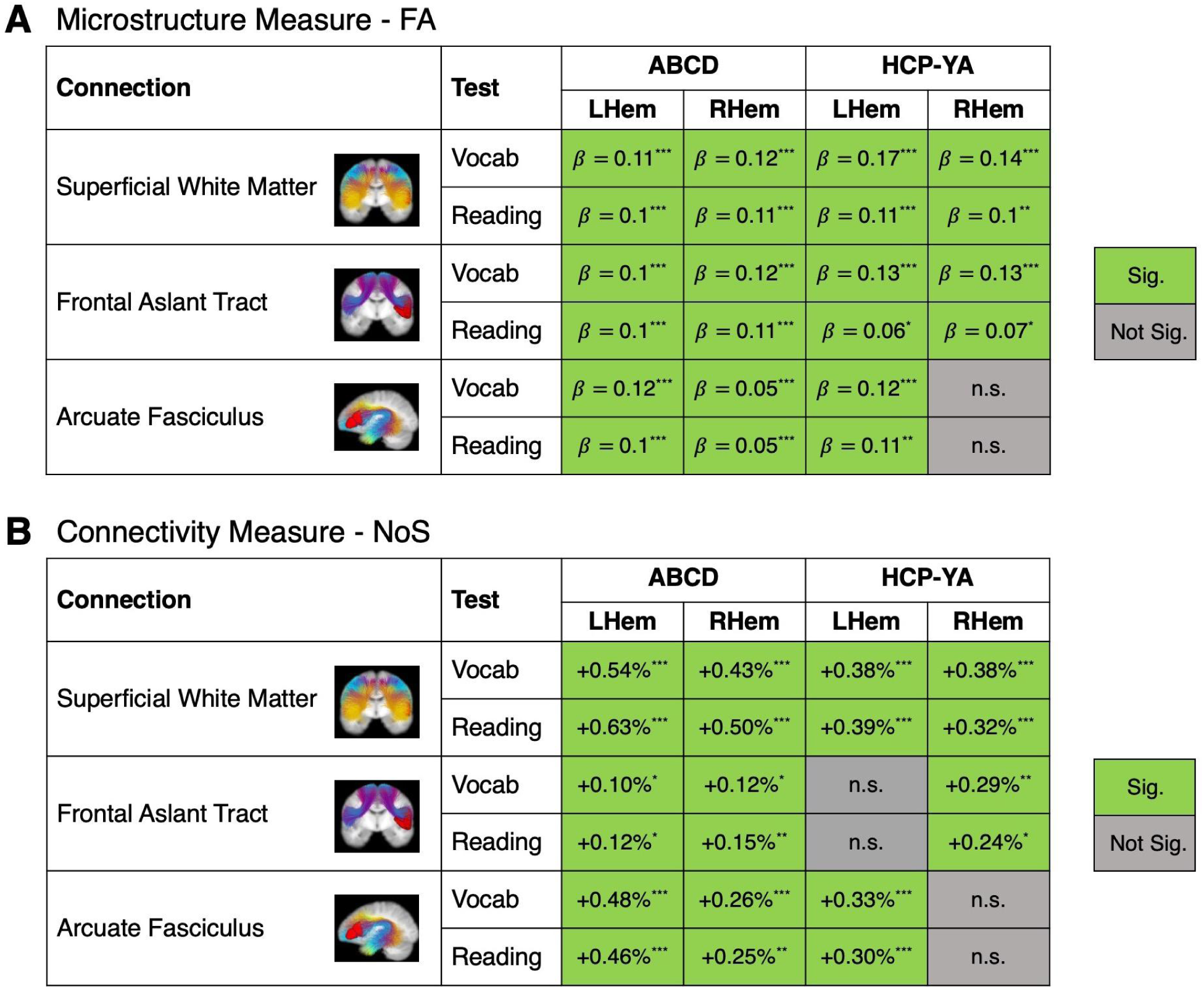
Broca’s area SWM microstructure (A) and connectivity (B) have significant associations with language performance (shown in green). The strength and direction of these relationships are quantified using standardized beta coefficients from linear regression models (A) and percent changes from negative binomial regression models (B). Significance coding (FDR-corrected): *** *p*<0.001, ** *p*<0.01, * *p*<0.05.

For the FAT, FA values are positively associated with language performance bilaterally in both age groups (all *p*<0.05). However, structural connectivity (NoS) of the FAT is bilaterally predictive of language performance in early adolescents (all *p*<0.05), but only in the right hemisphere in young adults (all *p*<0.05).

In the arcuate fasciculus, FA and NoS are positively associated with language performance bilaterally in early adolescents (all *p*<0.005). In young adults, FA and NoS of the arcuate fasciculus are positively associated with language performance exclusively in the left hemisphere (all *p*<0.005).

### 3.4 Structural lateralization of Broca’s area white matter connections relates to language performance

Finally, we examined whether structural lateralization influences language performance. Our analyses uncovered significant relationships between structural lateralization and language function, revealing pathway- and age-specific effects (Figure 8).

**Figure 8.**
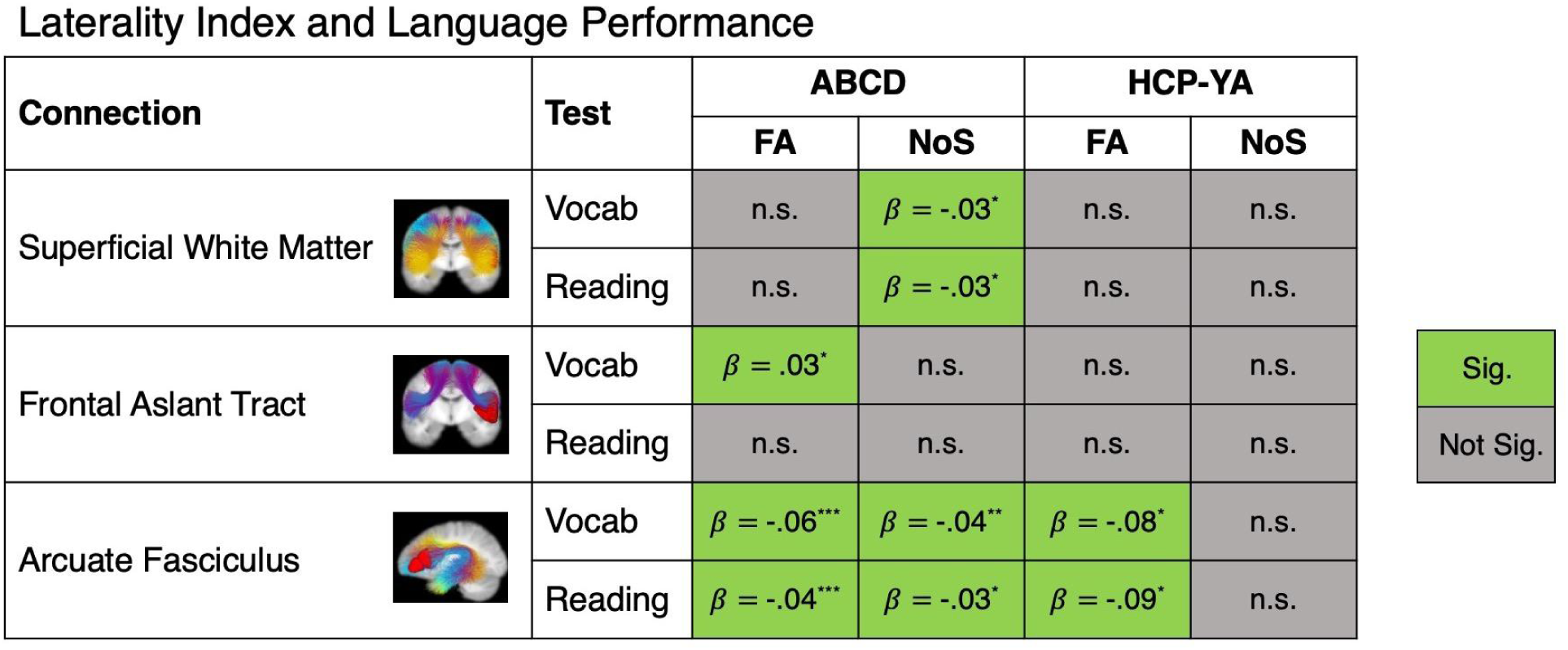
Structural lateralization of Broca’s area connections is associated with language performance. Relationships are quantified using standardized beta coefficients from linear regression models. Negative beta values indicate that greater leftward lateralization (LI) is associated with better language performance. Significance coding (FDR-corrected): *** *p*<0.001, ** *p*<0.01, * *p*<0.05.

For structural connectivity, multiple regression analyses showed that stronger leftward lateralization (LI of NoS) was associated with better language performance in early adolescents, specifically in Broca’s area SWM (*p*=0.02 for both PicVocab and ReadEng) and the arcuate fasciculus (*p*=0.006 for PicVocab and *p*=0.02 for ReadEng).

For microstructure, greater leftward microstructural lateralization (LI of FA) in the arcuate fasciculus was significantly linked to better language performance in both early adolescents (*p*<0.001 for PicVocab and ReadEng) and young adults (*p*=0.02 for PicVocab and ReadEng). The frontal aslant tract exhibited a contrasting pattern. In early adolescents, greater rightward microstructural lateralization (LI of FA) was significantly associated with better PicVocab performance (*p*=0.02).

## 4. Discussion

In this study, we characterized Broca’s area SWM and its relationship to individual language performance in two large, publicly available dMRI datasets, including over 10,000 participants spanning early adolescence and young adulthood. Overall, our findings demonstrate that Broca’s area SWM is significantly associated with language performance in both hemispheres in early adolescents and young adults. Notably, we observed that Broca’s area SWM exhibits greater hemispheric symmetry compared to the canonical arcuate fasciculus language pathway. Below, we contextualize these results with respect to existing literature and in the context of language function and neuroanatomy, highlighting key insights and implications.

Our findings both align with and extend prior research on the lateralization of SWM connections supporting language. An initial study examining frontal lobe connections did not identify any asymmetry of the short-range U-fibers between the middle frontal gyrus and superior frontal gyrus or between the inferior frontal gyrus and the middle frontal gyrus (Catani et al., 2012). A few initial studies have reported leftward asymmetries in the SWM using dMRI. For example, Phillips et al. (2013), using a white matter surface-based approach in 65 adults, identified leftward lateralization of FA in SWM across multiple areas, including the frontal lobe. Roman et al. (2022) observed left-lateralized quantitative anisotropy of frontal lobe SWM using a probabilistic tractography and fiber clustering framework in 100 HCP-YA participants. In line with this prior work demonstrating broad frontal lobe SWM microstructural lateralization, our focused study identified left-lateralized microstructure in Broca’s area SWM in both adolescents and young adults. In contrast to earlier studies that focused on microstructure, our analysis of Broca’s area SWM structural connectivity (i.e., NoS) revealed rightward lateralization in both adolescents and young adults. While this rightward structural lateralization has not been previously reported to our knowledge, other rightward structural lateralization is known in the frontal lobe, including the anterior segment of the arcuate fasciculus (Thiebaut de Schotten et al., 2011) and other branches of the superior longitudinal fasciculus (SLF) (Amemiya et al., 2021).

Overall, our results indicate a *dual lateralization pattern* of Broca’s area SWM, with significant leftward lateralization of FA and rightward lateralization of structural connectivity, in both adolescents and adults. This pattern is not contradictory; rather, it suggests an overall anatomical structure that is slightly “smaller” or less connected (in terms of NoS) in the left hemisphere, while its tissue architecture is more “orientation coherent” in terms of microstructure (FA). Our findings suggest the importance of both left and right hemisphere Broca’s area SWM for individual language function. However, the lateralization of Broca’s area SWM appears less important for language function, with the possible exception of adolescents, where greater leftward NoS lateralization (i.e., less of the observed rightward lateralization) was significantly associated with language functional performance, though with a small beta value.

These findings are consistent with studies of the lateralization of Broca’s area. Broca’s area is divided based on cytoarchitectonic criteria into a more posterior area generally corresponding to the pars opercularis, and a more anterior area 45 overlapping with the pars triangularis. Quantitative cytoarchitectonic studies generally find leftward asymmetry with larger volumes in left hemisphere area 44, but no asymmetry in area 45 (Amunts et al., 1999, 2003; Uylings et al., 2006).

For comparison, we examined FAT lateralization and found patterns consistent with prior studies (Broce et al., 2015; Catani et al., 2012; Linn et al., 2024; Pascual-Diaz et al., 2020; Szmuda et al., 2017; Taghvaei et al., 2024; Yeh, 2020). Although all laterality indices were statistically significant, the effect sizes in adolescents (for both FA and NoS) were very small, suggesting that significance may reflect our large sample size rather than strong hemispheric asymmetry in early adolescence. In adolescents, we observed slight but significant leftward FA and rightward NoS lateralization of the FAT, similar to Broce et al. (2015), who found nonsignificant leftward FA and significant rightward volume asymmetry in a small sample (n=19) of 5–8-year-olds. In contrast, in young adults, we observed more strongly left-lateralized FAT (FA and NoS), aligning with studies reporting leftward lateralization of various FAT properties, including volume, NoS, FA, and cortical coverage (Catani et al., 2012; Linn et al., 2024; Pascual-Diaz et al., 2020; Szmuda et al., 2017; Taghvaei et al., 2024; Yeh, 2020). Our findings also support bilateral relationships between FAT microstructure and language performance. Though limited literature exists on this topic, our findings are similar to recent work in the same HCP-YA young adult dataset, where left FAT cortical coverage was significantly associated with language scores, while right FAT FA showed a marginal association (Linn et al., 2024).

To contextualize our findings, we also examined the arcuate fasciculus as a benchmark for comparison with Broca’s area SWM. Leftward structural asymmetries of the arcuate fasciculus are well documented. For instance, Lebel & Beaulieu (2009) reported left-lateralized FA and NoS in the arcuate fasciculus in children and adults. Similarly, Broce et al. 2015 found left-lateralized microstructure in children, along with bilateral relationships between arcuate fasciculus microstructure and language performance. Sreedharan et al. 2015 also observed leftward lateralization of this pathway in children. Consistent with these findings, we observed left-lateralized microstructure and structural connectivity of the arcuate fasciculus in both early adolescents and young adults. Notably, we found that language performance was associated with bilateral arcuate fasciculus properties in adolescents, but exclusively with left hemisphere arcuate fasciculus properties in adults.

Compared to the SWM and FAT, the arcuate fasciculus exhibited the greatest degree of leftward lateralization, particularly in terms of structural connectivity (NoS). The laterality indices for the SWM and FAT were notably smaller in magnitude than those of the arcuate fasciculus, indicating more symmetric structural organization in adolescents and young adults. The comparatively symmetric lateralization observed in the SWM and FAT suggests that these more superficial or frontally localized pathways may support language in a more bilaterally distributed manner. Although the arcuate fasciculus showed the most pronounced leftward structural lateralization, its association with language performance was not stronger than that of the less-lateralized SWM or FAT. This suggests that structural lateralization does not necessarily imply greater functional involvement. This structure-function dissociation aligns with prior findings indicating that anatomical and functional lateralization do not always correspond directly (Ocklenburg & Güntürkün, 2024; Vernooij et al., 2007).

Our findings suggest stronger left hemisphere microstructural (FA) lateralization of language-related white matter with age. These findings align with previous work reporting age-related increases in left hemisphere lateralization in canonical language pathways, such as the arcuate fasciculus (Broce et al., 2015; Tak et al., 2016), as well as findings from quantitative human cytoarchitectonic analyses showing increased left lateralization with age (Amunts et al., 2003). While functional MRI studies in adults have demonstrated pronounced leftward lateralization of functional language networks, findings in children suggest a more bilateral organization of these networks (Gaillard et al., 2000; Knecht, Deppe, et al., 2000; Knecht, Dräger, et al., 2000; Szaflarski et al., 2002, 2006). The structural underpinnings of this functional lateralization, particularly in white matter connections to Broca’s area, remain an area of active investigation.

The present results fill a gap in the understanding of the connectional neuroanatomy of Broca’s area and highlight the complexity of understanding the connectivity of brain areas involved in language in the human brain. The direct visualization of structural neural pathways, from origin through pathway to termination, is only possible in experimental animals (Rushmore et al., 2020). The macaque monkey is the closest experimental model of the human brain in which large-scale connectional analyses based on tract tracers have been performed (e.g., (Bakker et al., 2012; Felleman & Van Essen, 1991; Yeterian et al., 2012)). However, results from the macaque model have limited applicability for modeling language-related regions and networks, given differences in language-related functions between the two species, and because tracer studies rarely evaluate differences between hemispheres. Thus, even though considerable information is available in the monkey regarding the complex connectivity of frontal lobe regions homologous to Broca’s area (e.g., (Barbas & Pandya, 1989; Frey et al., 2014; Petrides & Pandya, 2006, 2009; Yeterian et al., 2012), the current study provides critical data about the connectivity of Broca’s area that cannot be obtained with experimental animal models.

Our findings highlight the potential importance of superficial white matter (SWM) in neurosurgical planning (Latini & Ryttlefors, 2015), where preserving language function is critical (Essayed et al., 2017; Silva et al., 2018). Mapping these SWM connections in individual patients (Xue et al., 2023), especially in conjunction with personalized mapping of language-associated cortex, could be useful for guiding surgical planning. Further study of the finer subdivisions within the SWM is also warranted, as their functional roles may differ, and surgical interventions typically affect only a portion of the broader SWM region examined here.

Finally, we note some limitations of this study and directions for future research. First, this study focused on the superficial white matter of Broca’s area, a critical language region; however, this provides only a partial view of the broader language network. Future work could examine superficial white matter connectivity involving other key regions, including temporoparietal areas, to gain a more comprehensive understanding of the white matter architecture supporting language. Second, although our study of early adolescents and young adults may offer insight into developmental differences, the cross-sectional nature of the data limits direct inference about within-subject changes over time. Longitudinal studies are needed to more precisely characterize developmental trajectories in language-related white matter. Third, we investigated SWM lateralization in the general population; future work may consider analyses by handedness group (Taghvaei et al., 2024) or handedness degree (Propper et al., 2010). Fourth, there are inherent differences between the ABCD and HCP-YA datasets, including sample size and acquisition parameters such as spatial resolution. While we observed notable consistencies in our findings across both datasets, these methodological differences may contribute to variability and should be considered when interpreting cross-dataset comparisons. Finally, while this study focused on Broca’s area SWM as a whole, we demonstrated its fine-grained parcellation into multiple fiber clusters. Future work is needed to examine the lateralization of these individual fiber clusters and their specific relationships to language function.

## 5. Conclusion

In conclusion, this study provides a large-scale investigation of Broca’s area SWM and its relationship to language performance in early adolescents and young adults. By analyzing over 10,000 participants, we find that Broca’s area SWM exhibits distinct patterns of structural lateralization and is bilaterally associated with language performance in both adolescents and young adults. Compared to the more strongly lateralized arcuate fasciculus, Broca’s area SWM exhibits more bilateral structural organization and relationships with language performance. Our findings suggest that short-range and medium-range frontal pathways, including the SWM and FAT, may provide bilateral support for language processing from adolescence into adulthood. Our results contribute to an evolving model of the neurobiology of language that extends beyond classical left-lateralized networks to encompass a broader, more distributed white matter architecture and underscore the importance of considering the SWM in both developmental research and clinical applications, such as neurosurgical planning.

## Data and Code Availability

The data used in this project are from the Adolescent and Cognitive Development Study (ABCD) dataset, accessible via the National Institute of Mental Health Data Archive (NDA) at <u>nda.nih.gov/abcd</u>, and the Human Connectome Project Young Adult (HCP-YA) dataset, available through the ConnectomeDB at db.humanconnectome.org. Additionally, we leveraged the harmonized baseline ABCD data, available via the NDA (see (Cetin-Karayumak et al., 2024)). The ORG atlas and associated code for applying the atlas are publicly available at https://dmri.slicer.org/atlases/.

## Author Contributions

Shiva Hassanzadeh-Behbahani: Conceptualization, Methodology, Formal Analysis, Visualization, Writing - Original Draft. Zhou Lan: Formal analysis, Methodology, Writing - Review & Editing. Leo Zekelman: Resources, Methodology, Writing - Review & Editing. R. Jarrett Rushmore: Funding Acquisition, Methodology, Writing - Review & Editing. Yuqian Chen: Software, Writing - Review & Editing. Tengfei Xue: Software, Writing - Review & Editing. Suheyla Cetin-Karayumak: Software, Writing - Review & Editing. Steve Pieper: Software, Writing - Review & Editing. Yanmei Tie: Writing - Review & Editing. Edward Yeterian: Methodology, Writing - Review & Editing. Alexandra J. Golby: Funding Acquisition, Writing - Review & Editing. Nikos Makris: Methodology, Writing - Review & Editing. Fan Zhang: Software, Writing - Review & Editing. Yogesh Rathi: Software, Writing - Review & Editing. Lauren J. O’Donnell: Conceptualization, Methodology, Funding Acquisition, Software, Supervision, Writing - Review & Editing.

## Acknowledgments

Data used in the preparation of this article were obtained from the Adolescent Brain Cognitive Development (ABCD) Study (https://abcdstudy.org), held in the NIMH Data Archive (NDA). This is a multisite, longitudinal study designed to recruit more than 10,000 children aged 9-10 and follow them over 10 years into early adulthood. The ABCD Study® is supported by the National Institutes of Health and additional federal partners under award numbers U01DA041048, U01DA050989, U01DA051016, U01DA041022, U01DA051018, U01DA051037, U01DA050987, U01DA041174, U01DA041106, U01DA041117, U01DA041028, U01DA041134, U01DA050988, U01DA051039, U01DA041156, U01DA041025, U01DA041120, U01DA051038, U01DA041148, U01DA041093, U01DA041089, U24DA041123, U24DA041147. A full list of supporters is available at https://abcdstudy.org/federal-partners.html. A listing of participating sites and a complete listing of the study investigators can be found at https://abcdstudy.org/consortium_members/. ABCD consortium investigators designed and implemented the study and/or provided data but did not necessarily participate in the analysis or writing of this report. This manuscript reflects the views of the authors and may not reflect the opinions or views of the NIH or ABCD consortium investigators. The ABCD data repository grows and changes over time. The ABCD data used in this report came from DOI: 10.15154/1520591 (Data Release 3.0).

Human Connectome Project Young Adult (HCP-YA) data were provided, in part, by the Human Connectome Project, WU-Minn Consortium (Principal Investigators: David Van Essen and Kamil Ugurbil; 1U54MH091657) funded by the 16 NIH Institutes and Centers that support the NIH Blueprint for Neuroscience Research; and by the McDonnell Center for Systems Neuroscience at Washington University.

## Footnotes

1 github.com/pnlbwh/ukftractography

2 https://github.com/SlicerDMRI/whitematteranalysis

