## Supplementary Table 1 for "The superficial white matter in language processing: Broca’s area connections are bilaterally associated with individual performance in children and adults"

### Supplementary Material

**Supplementary Table 1.** Fiber cluster indices from Broca's area white matter in the ORG Atlas.

| Connection | ORG Atlas Cluster Numbers |
|---|---|
| Arcuate Fasciculus | 169, 170, 171, 176, 181, 186, 206, 726 |
| Frontal Aslant Tract | 267, 299, 300, 322, 361, 375, 377 |
| Superficial White Matter | 205, 262, 276, 298, 359, 373, 591, 593, 603, 607, 635, 639, 645, 762 |
